## Supplementary Material for "Intrinsic macroscale oscillatory modes driving long range functional connectivity detected with ultrafast fMRI"

**This file includes:**

Supplementary Text
Figures S1 to S17
Tables S1 to S2

**Other Supplementary Materials for this manuscript include the following:**

Video S1

Caption:
**Video S1 - fMRI signals band-pass filtered in the frequency range that best differentiated between conditions in 3 different rats and in 3 different conditions.** After removing the mean from the fMRI signals in each voxel, the signals were bandpass filtered between 0.15 and 0.25 Hz and are imaged over time. Sedation: Medetomidine only; Light Anesthesia: Medetomidine + 1% isoflurane; Deep anesthesia: Medetomidine + 3% isoflurane). To account for expected differences in power across conditions, colorbar limits are set to ± 4 standard deviations of the band-pass filtered signals in each scan.

### Supplementary Text

I. Space-frequency analysis up to the Nyquist frequency

The unprecedented time resolution of our acquisitions allows performing a space-frequency analysis up to the Nyquist frequency of 13 Hz. As shown in Fig. S5, power >0.4 Hz in fMRI signals above postmortem levels was found to relate to non-cortical physiological rhythms such as:

i) Breathing: A clear frequency peak was detected between 0.8 and 1.7Hz in all scans, matching the values recorded experimentally (green patch in Fig. S5 C; breathing frequency reported for each scan in Table S1). Notably, power at the breathing frequency was detected mostly in non-cortical voxels.

ii) Heartbeat: A clear frequency peak was detected in all scans between 4 Hz and 6Hz resonating significantly above postmortem levels in non-cortical voxels in the expected location of the main arteries.

iii) Harmonics of physiological rhythms: Peaks at twice the breathing and heartbeat frequencies were detected in all scans.

Additionally, a resonant frequency at 7.6 Hz associated to scanner noise was detected in all scans including the postmortem scans (see black dotted line in Fig. S5).

No other frequencies >0.4Hz were found to resonate above chance levels with the current resolution.

On the other hand, power in brain voxels was found to peak below 0.4 Hz (Fig. S5 BC). Voxel-wise power spectra were re-calculated with 5000 frequency bins of 0.0053 Hz each, and spectral maps were obtained for 0.05Hz-wide frequency bands <0.5 Hz Fig. S5 D). Increasing the frequency specificity reveals a frequency range where strong power is detected in cortical voxels.

II. Effect of the temporal resolution in fMRI scans

We evaluated the sensitivity of our results to the temporal resolution of fMRI acquisitions, by repeating the analysis after down-sampling the fMRI data, i.e. by taking the signal every 2 frames (equivalent to TR=76 ms), every 3 frames (TR=114 ms), and so on. Focusing on the narrow frequency band that maximally differentiated between conditions (i.e., 0.20-0.24 Hz) we verified that the power in this band decreases with increasing TR, compromising the detection of significant differences in the mean power between conditions (Fig. S13).

III. Analysis of whole-brain fMRI scans

Prior to the analysis of resonant modes, the multi-slice fMRI acquisitions from 3 different rats were realigned and corrected for slice timing using linear interpolation. Following a spectral analysis of each individual scan up to the Nyquist frequency of 1.4 Hz, the fMRI signals from brain voxels were band-pass filtered between 0.05 and 0.25Hz to remove artefacts detected below 0.05 Hz and > 0.25Hz (Fig. S11). The covariance matrices were computed for each scan in this frequency range and were averaged across scans. Following the same procedure of the single slice analysis, the eigenvectors of the average covariance matrix were extracted. Since no postmortem scan was performed to define a baseline threshold for the associated eigenvalues, the 15 eigenvectors with largest magnitude eigenvalues were considered in the subsequent analysis (Fig. S10).

### IV. Data and Code availability

All MRI data, code and materials used in this study are available in https://drive.google.com/drive/folders/1s99k5H1riQV-QONHrn-6cEphbv2VctLv?usp=sharing, which will be deposited in a public database upon acceptance.
The data is organized in the following folders:

**MRI data:** Folder Manuscript_MRI_data contains the single-slice (sub-folder Frontal_Slice) and multi-slice (sub-folder WholeBrain) recordings already imported from the scanner to .mat format (using code Import_scans_server.m). Each .mat file contains the images of a given scan in 2D (anatomical single slice), 3D (functional single-slice) or 4D (functional multi-slice), together with the methods and acquisition parameters.

**Analysis codes:** Codes for the MRI data analysis and generation of all the figures of the manuscript are made available in folder Manuscript_Codes. The first code is fMRI_SpectralMaps.m, which performs the space-frequency analysis saving the power maps in each frequency band, as shown in Figure 1A and Supplementary Figures 3, 4, 5 and 7). Parameters for realignment and the brain masks are previously saved (but can be replicated using function DefineMasks.m). Subsequently, code Modes_in_bands.m computes the eigenvectors of the covariance matrices in each band, and Wave_modes.m calculates the power spectrum and the resonance Q-factor for each mode. Figure 1 is generated with code FC_seed_correlation.m Figure 2 is generated with Figure_SpecMap_Bands.m, Figures 3 and 4 with Plot_wave_modes.m, and Figure 3 with Model_Sum_Waves.m. Simulation plots in Figure 4 are generated using Hopf_One_unit_fMRI.m.

**Figures and Videos:** Original figure files in .fig and .pptx are provided in folder Manuscript_Figures. High resolution versions of the videos included as supplementary files are provided in folder Manuscript_Videos.

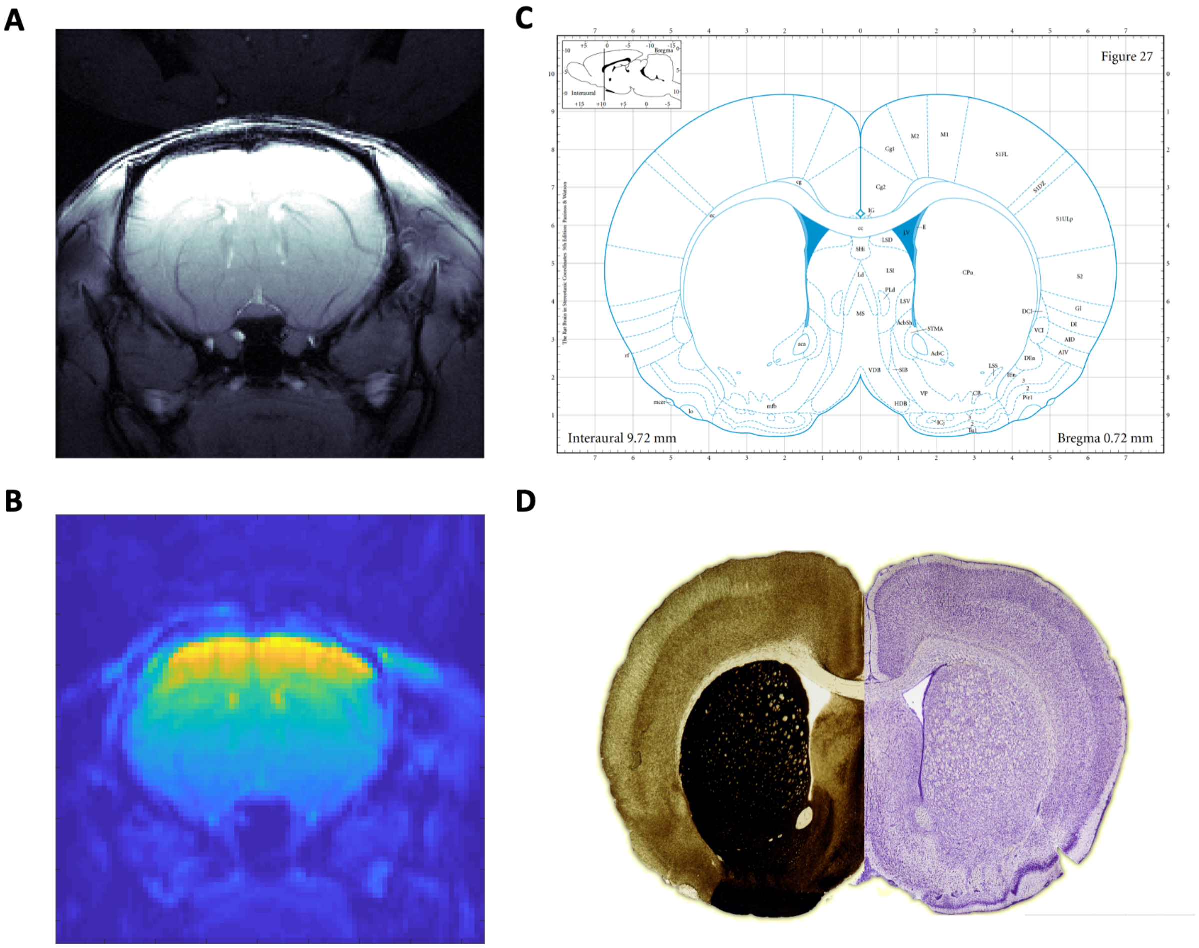


**Fig. S1. Frontal slice selected for ultrafast resting-state fMRI recordings of the rat brain.** (A) Multi gradient echo (MGE) anatomical image (TE = 2.5:5:97.5 ms) from a representative rat. (B) Average fMRI image (GE-EPI) obtained from the same rat. (C) Anatomical diagram from the Paxinos & Watson rat brain atlas. (D) Plate image using different staining methods (left – AchE; right – cresyl violet).

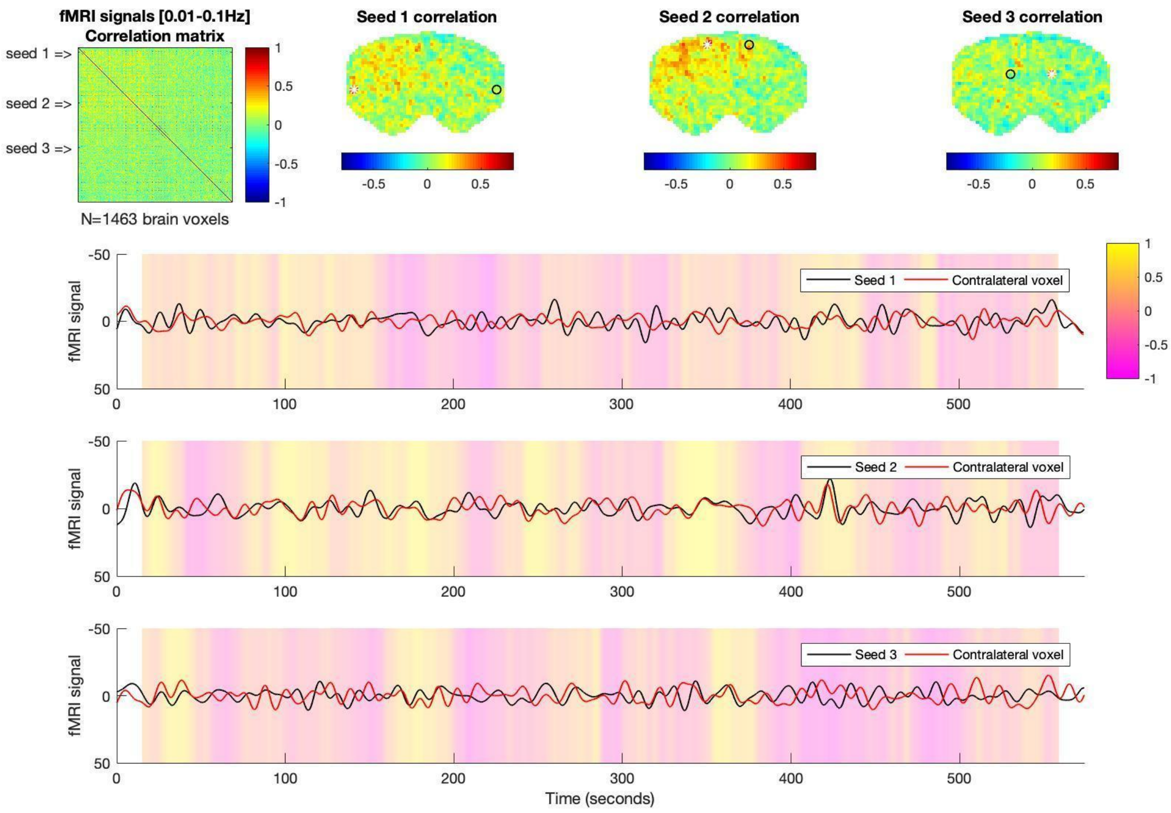


**Fig. S2 – Resting-state functional connectivity analysis in postmortem scan.** (a) Correlation matrix of the fMRI signals in all voxels within the brain mask, bandpass filtered in the range typically considered in resting-state studies, i.e., 0.01-0.1Hz (no nuisance regressor nor spatial smoothing applied). Each line/column in the matrix corresponds to the correlation map of each voxel. (b) Seed-based correlation maps are represented for 3 different seeds (white asterisks), where each voxel is colored according to its degree of correlation with the seed. A voxel contralateral to each seed is represented by a black circle. All colorbars are truncated between -0.8 and 0.8. (c) Filtered fMRI signals recorded in each seed (red) and corresponding contralateral voxel (black). Colored shades represent the sliding window correlation (SWC) using a 30-second window. Although periods of positive or negative SWC can still be detected, these last visibly shorter and, when considering the entire recording time, no particular pattern of long-range connectivity is detected (except from a subtle left-right gradient associated with the MRI acquisition).

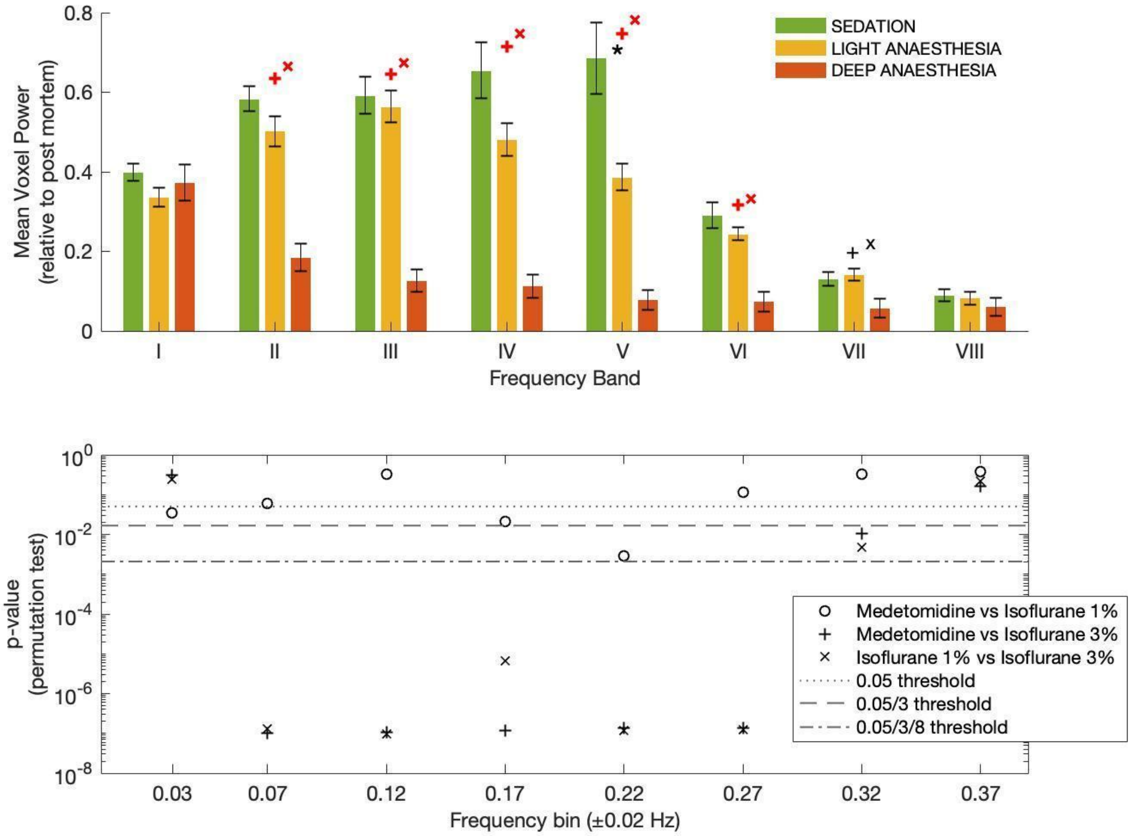


**Fig. S3. Statistical significance of differences in spectral power between conditions.** (Top) Mean voxel power in each frequency band where zero corresponds to the mean power in postmortem scans. Error bars report the mean ± standard error across scans in each condition. Symbols indicate statistical significance between: ‘o’ sedation vs light anaesthesia; ’+’ sedation vs deep anaesthesia; ‘x’ light vs deep anaesthesia, corrected by the number of conditions (black $p<0.0167$), or by the number of conditions*bands (red, $p< 0.0021$). (Bottom) The p-values of the permutation test, showing their position with respect to the different threshold lines. The spectral power is found to vary significantly and nonlinearly across conditions.

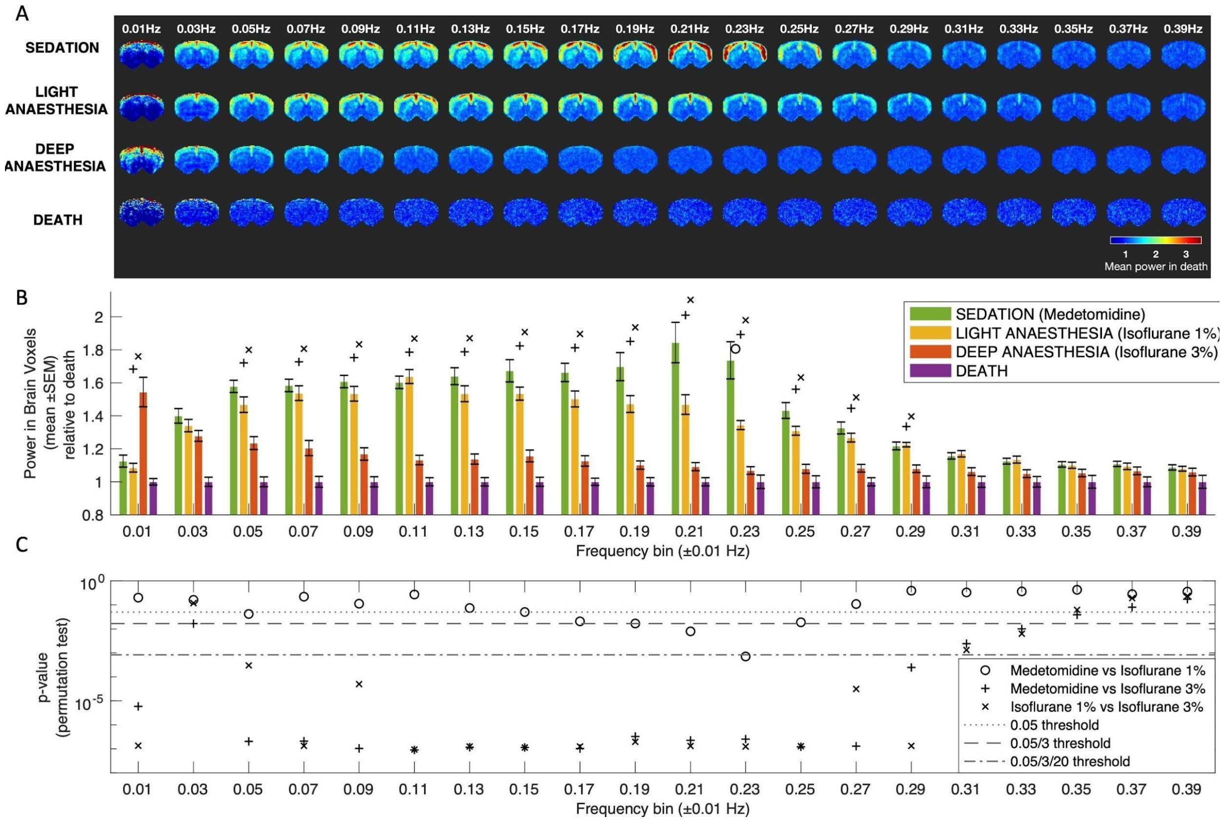


**Fig. S4. Space-frequency analysis at narrower frequency bands.** (A) Average power across scans in the same condition calculated in each frequency band (using frequency bins of 0.02Hz). (B) The total power in brain voxels was statistically compared between scans in the different conditions in each frequency range. (C) The *p*-values of the statistical comparison with respect to conservative thresholds correcting for multiple comparisons considering all the comparisons performed, here across 20 distinct frequency bands. All frequency bins up to 0.30 Hz (except one centered at 0.03Hz) were found to differentiate significantly between at least 2 conditions. On one side, the brains of sedated rats were found to consistently exhibit relatively high power in all bins between 0.02 and 0.30 Hz, peaking at 0.21 Hz. Under light anaesthesia however, this peak is significantly decreased, and the strongest power is detected instead around 0.11Hz. Deep anesthesia was found to significantly change the power in all frequency bins up to 0.30 Hz ($p<0.00083$ surviving Bonferroni correction for multiple comparisons, see Supplementary Figure 6BC). On one hand, oscillations between 0.04 and 0.30 Hz were found to decrease in power with 3% isoflurane. On the other, oscillations below 0.02 Hz were found to increase concomitantly under deep anaesthesia.

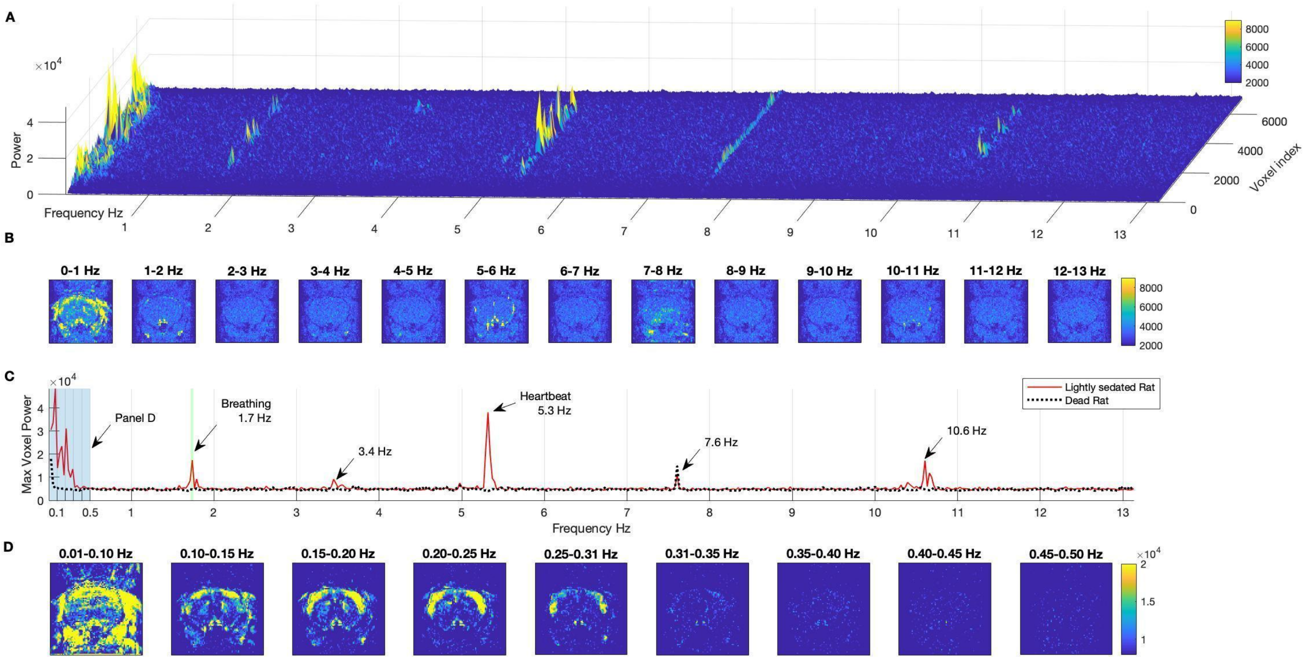


**Fig. S5. Analysis of fMRI spectral content up to Nyquist Frequency.** (A) Power spectra of the fMRI signal in each of the 7056 voxels of a frontal brain slice of a rat under light sedation (medetomidine). A number of resonant frequencies is detected, as highlighted by yellow peaks, showing increased power at relatively narrow frequency ranges and for specific voxels. (B) For a range of frequency bands covering the whole spectrum, voxels exhibiting increased power at the corresponding frequency are highlighted in yellow (using the same colour scale as (A)). (C) Power spectrum showing the maximum power detected across voxels (to highlight emerging rhythms irrespective of the number of voxels exhibiting it). A peak matching the experimentally recorded breathing frequency (green patch) was detected. Harmonics were detected at twice the breathing (1.7 Hz x 2 = 3.4 Hz) and heartbeat (5.3 Hz x 2 = 10.6 Hz) frequencies. A peak at 7.6 Hz relating to the MRI sequence was detected both in the live and dead animal. (D) Power above noise levels was only detected in brain voxels for frequencies below 0.5 Hz, revealing the intrinsically slow nature of spontaneous brain activity, with clear power detected within cortical boundaries particularly between 0.20 and 0.25 Hz.

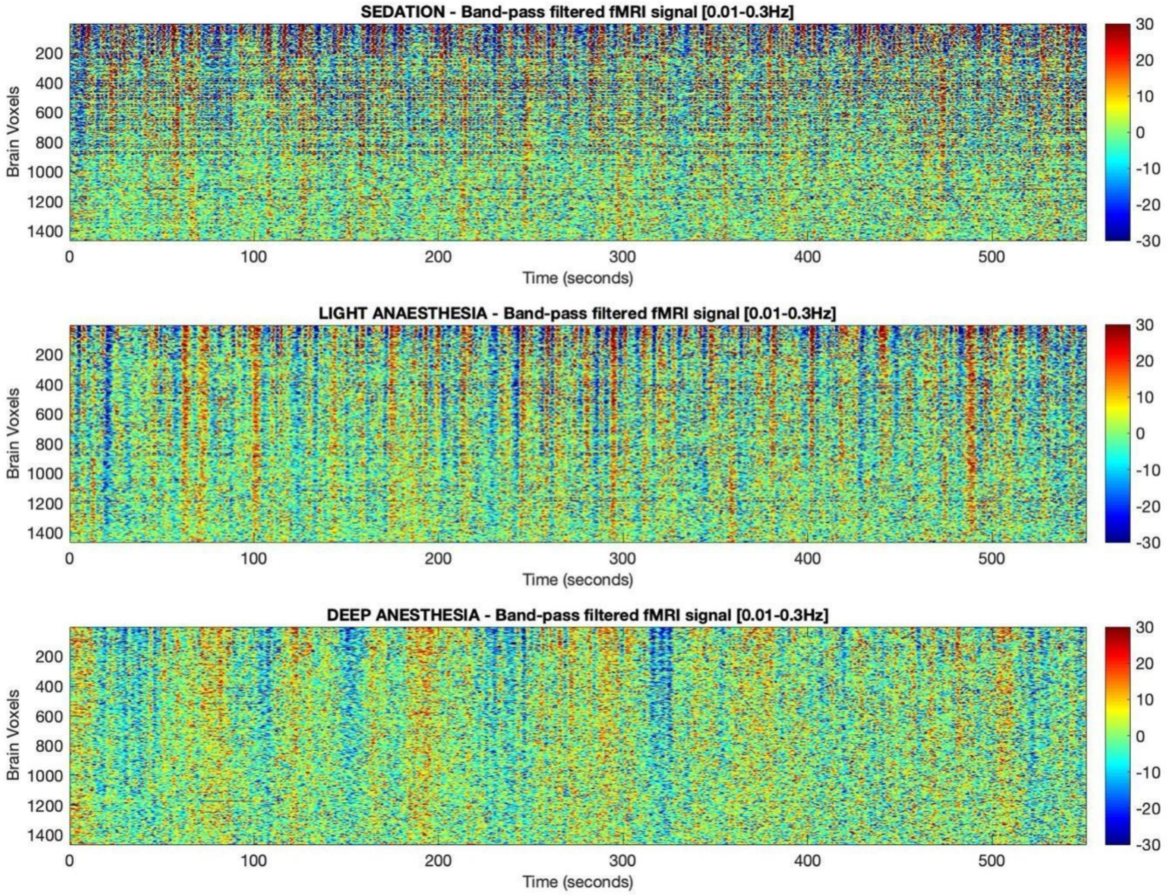


**Fig. S6. Carpet plot showing the fMRI signals from the same rat in 3 experimental conditions**. The fMRI signals in each voxel and each time point (1463 voxels x 16000 time points) are plotted after applying band-pass filters between 0.01 and 0.3Hz, covering the range where the oscillatory power in the brain was significantly above postmortem levels. fMRI signals from rat 1 under (Top) sedation, medetomidine only), (Middle) light anesthesia, medetomidine + 1% isoflurane and (Bottom) deep anesthesia, medetomidine + 3% isoflurane. Signals exhibit distinct spatiotemporal properties across conditions.

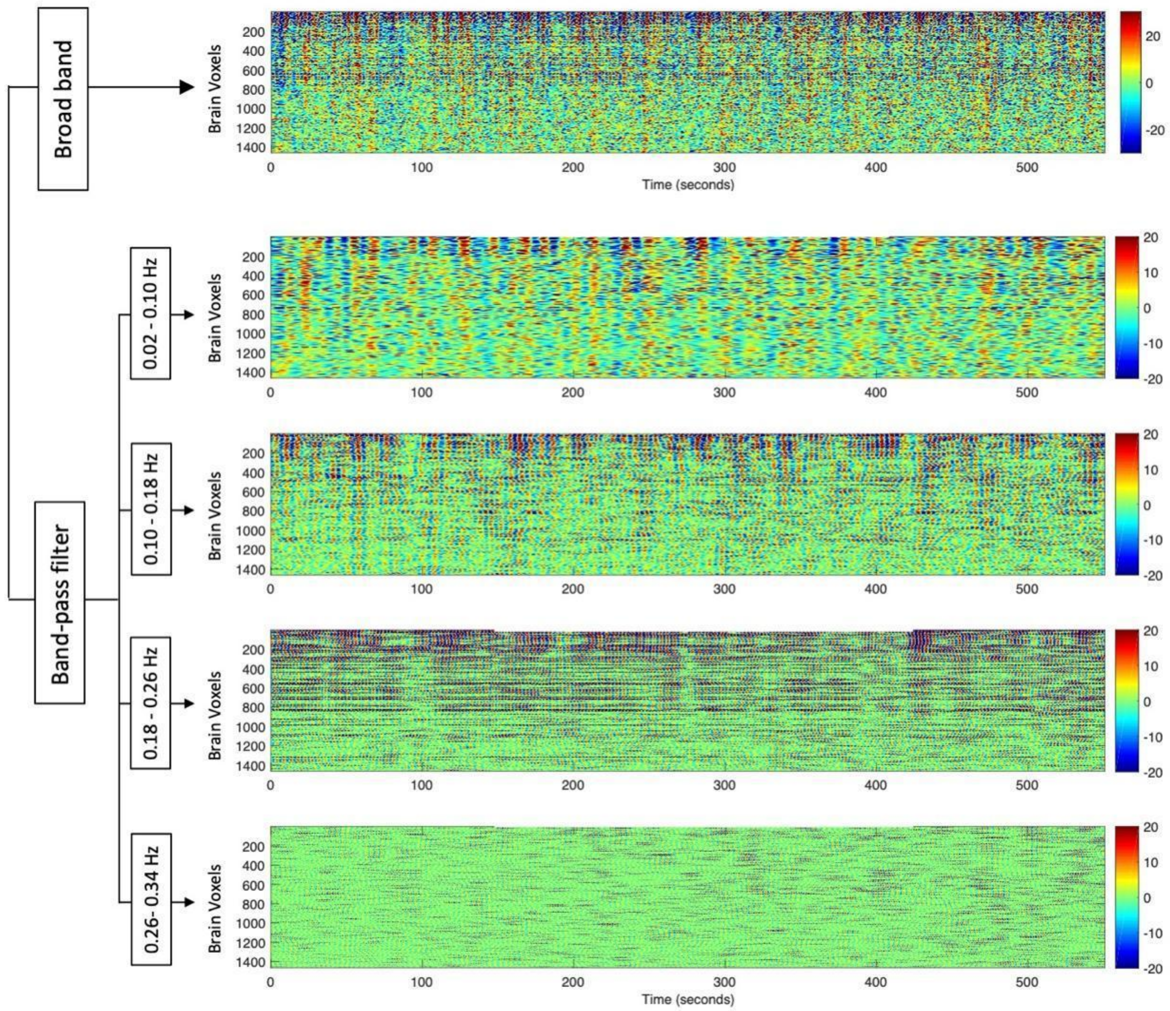


**Fig. S7. Carpet plot showing the fMRI signals from a representative scan of a sedated rat.** The fMRI signals from rat 1 under medetomidine only are plotted for each voxel and each time point (1463 voxels x 16000 time points), unfiltered (Top) and after applying band-pass filters in 4 non-overlapping frequency bands covering the range where the oscillatory power in the brain was significantly above postmortem levels. Sustained oscillatory patterns are found to emerge transiently at different frequency bands and in different voxels.

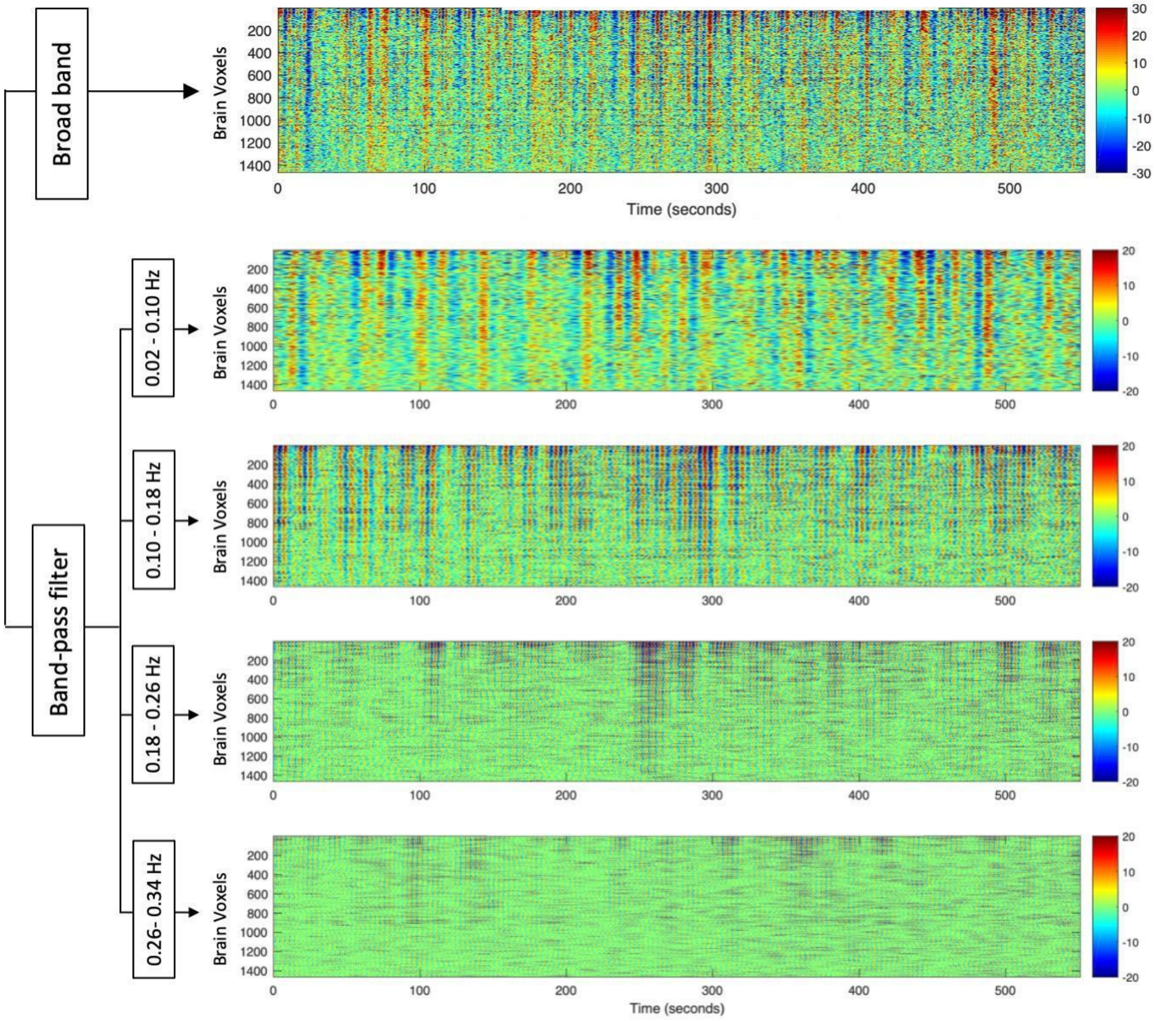


**Fig. S8. Carpet plot showing the fMRI signals from a representative scan of a lightly anesthetized rat.** The fMRI signals from rat 1 under medetomidine + 1% isoflurane are plotted for each voxel and each time point (1463 voxels x 16000 time points), unfiltered (Top) and after applying band-pass filters in 4 non-overlapping frequency bands covering the range where the oscillatory power in the brain was significantly above postmortem levels. Sustained oscillatory patterns are found to emerge transiently at different frequency bands and in different voxels. Compared to the isoflurane-free condition, sustained oscillatory patterns are found to be more globally synchronized across voxels, and display visibly lower power in the 0.18-0.26 Hz range.

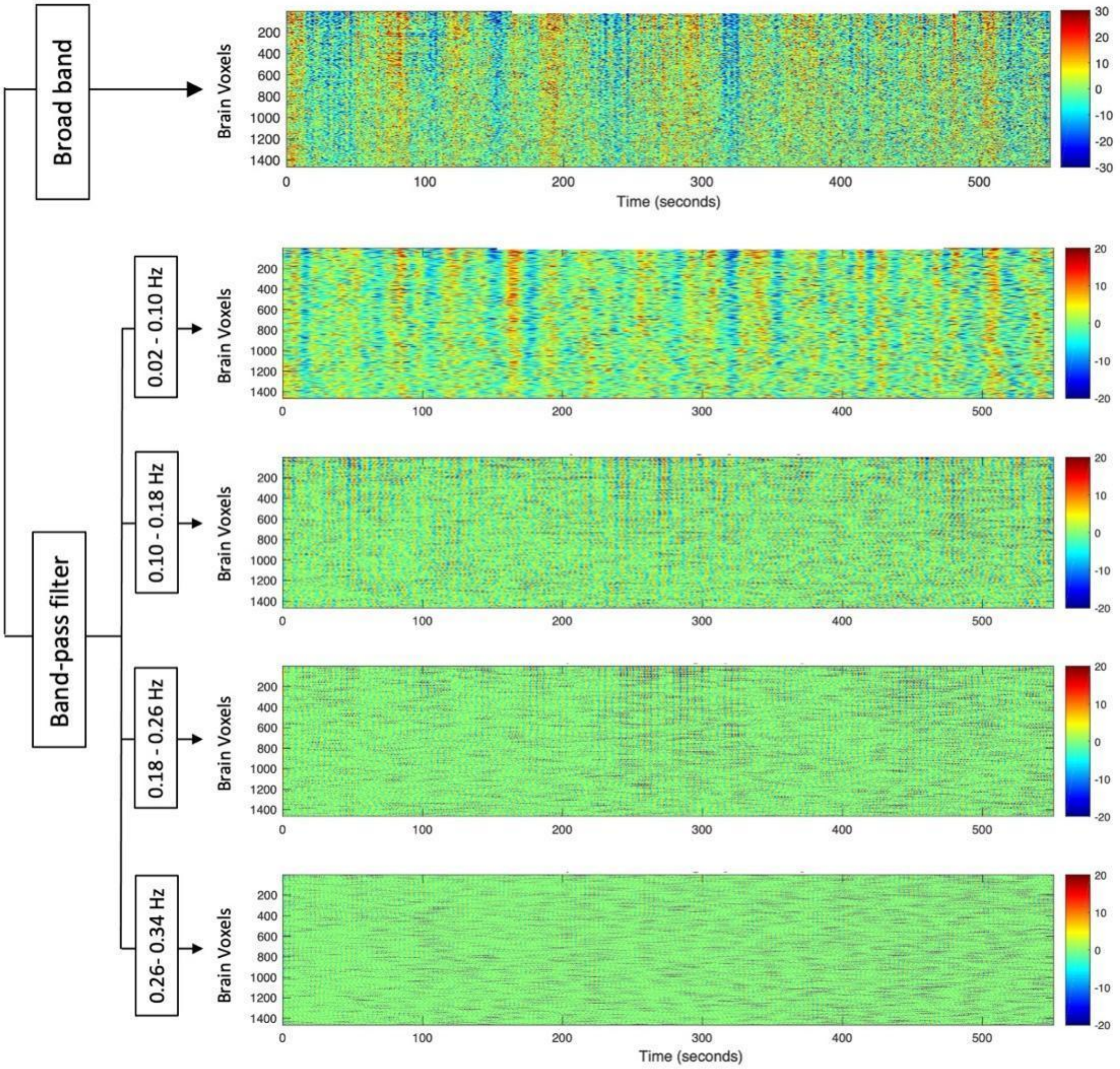


**Fig. S9. Carpet plot showing the fMRI signals from a representative scan of a deeply anaesthetized rat.** The fMRI signals from rat 1 under medetomidine + 3 % isoflurane are plotted for each voxel and each time point (1463 voxels x 16000 time points), unfiltered (Top) and after applying band-pass filters in 4 non-overlapping frequency bands covering the range where the oscillatory power in the brain was significantly above postmortem levels. Sustained oscillatory patterns are found to emerge transiently at different frequency bands and in different voxels. Under deep anaesthesia, the activity is dominated by slower, less periodic and more global patterns, with visibly lower power> 0.1 Hz.

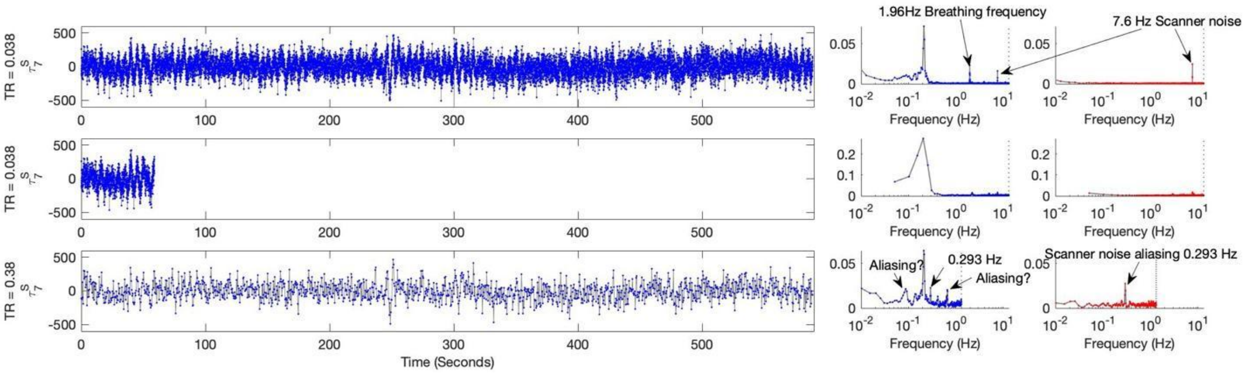


**Fig. S10 - Effect of sampling rate and scan duration on the power spectrum of the temporal signature associated with a principal component.** (Top) The raw TxN fMRI signals with N=1463 voxels and T=15501 time points are multiplied by the Nx1 principal component, resulting in a 1xT time series. The associated power spectrum is shown on the right (blue), with frequency resolution of 0.0064Hz and a Nyquist frequency of 13.1579 Hz (vertical dashed line) using the default parameters of Welch's power spectral density estimate in Matlab. The frequency axis is in log scale to highlight both low and high frequency components. On the far right panel, the power spectrum is reported for a time series obtained by multiplying the same principal component to the raw signals of a postmortem scan (red). (Middle) The same analysis is performed by considering the same sampling rate but only 1/10th of the total scan duration, resulting in 1551 time points. With reduced scan duration, the Nyquist frequency is still 13.1579 Hz but the frequency resolution is reduced to 0.0514 Hz. (Bottom) The same analysis is performed by reducing the sampling rate by 1/10th while keeping the original scan duration, resulting also in 1551 time points. With reduced sampling rate, the Nyquist frequency is 1.3158 Hz and the frequency resolution is 0.0051 Hz. This analysis shows that while long scans with low sampling rate may have the same (or even higher) frequency resolution as the original fast sampled signal, the undersampled signals can exhibit artifacts resulting from aliasing of high frequency components in the signal both from physiological sources (breathing and heartbeat) or from scanner artifacts (present also in postmortem scans).

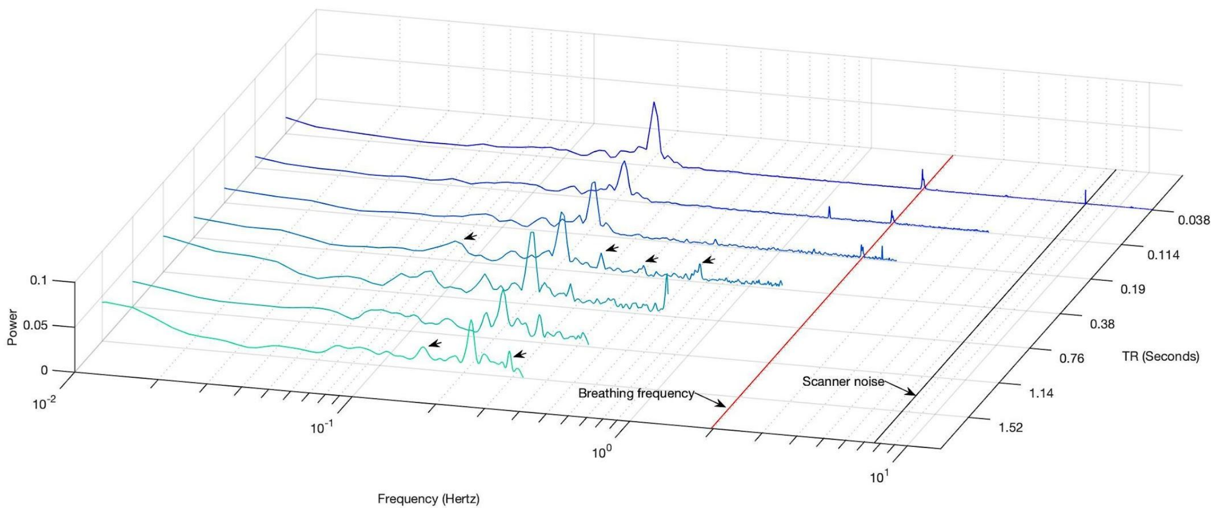


**Fig. S11 - Effect of the sampling rate in frequency aliasing.** The power spectra are calculated for the same signal (fixed scan duration of 10 minutes) with different downsampling factors S=1,3,5,10,20,30,40 ranging from the original Repetition Time (TR) of 38 ms (darker blue) to 1.52 seconds (by considering only one in every S=40 images) (cyan). As the downsampling S is increased, the Nyquist frequency is reduced as fNyq=1/(2*S*TR). When the Nyquist frequency is below the breathing frequency (red line), the breathing frequency cannot be adequately resolved, and new frequency peaks appear in the power spectrum (see the black arrows for TR=380 ms).

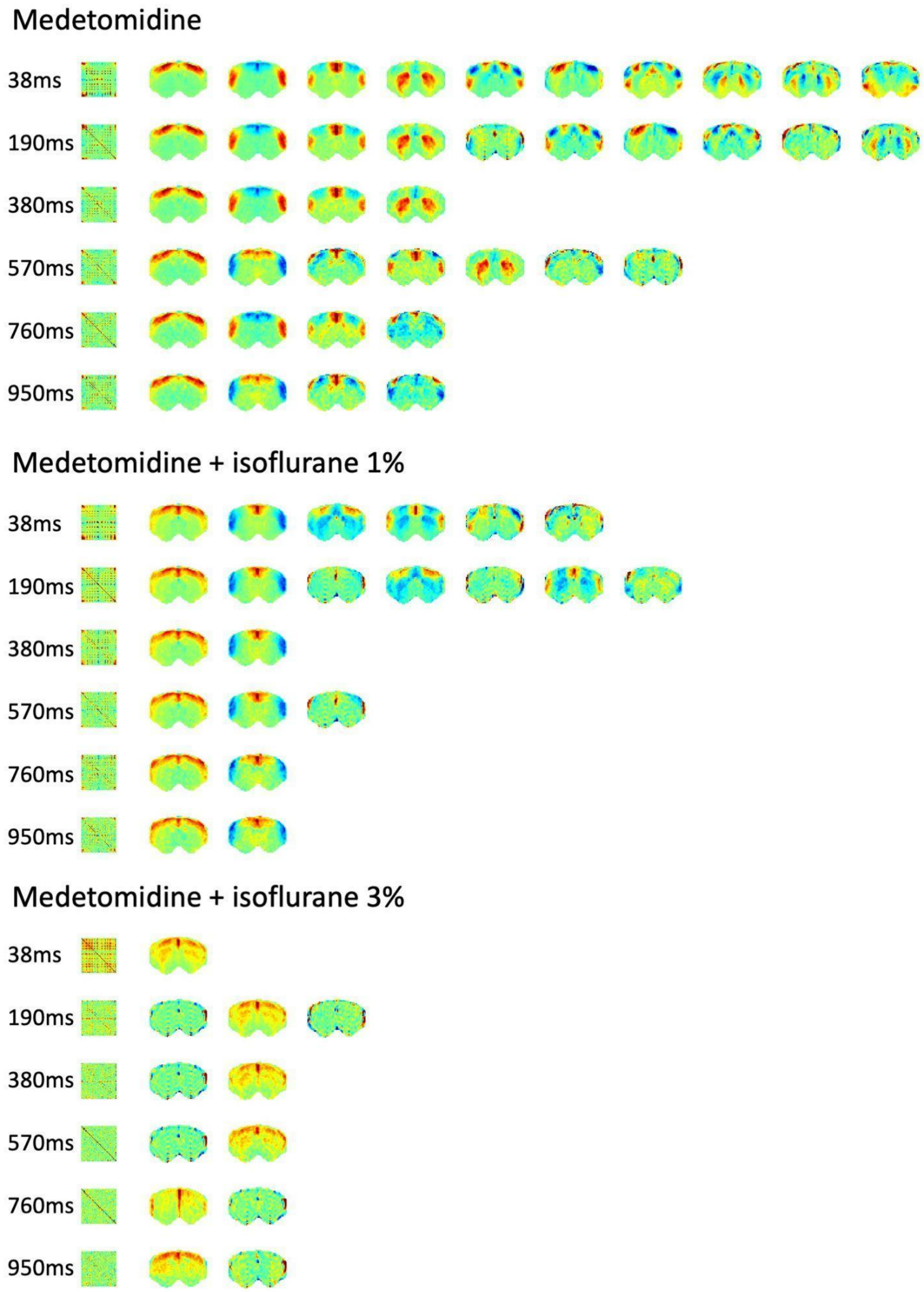


**Fig. S12 - Effect of the sampling rate on the number and spatial signature of the principal components detected.** Before the analysis, the fMRI signals recorded with a TR of 38 milliseconds (ms) were downsampled by considering only one in every 5, 10, 15, 20 and 25 frames (corresponding to intervals of 190, 380, 570, 760 and 950 milliseconds between frames). The number of principal components detected corresponds to the number of eigenvalues larger than the largest eigenvalue from postmortem scans (signals also downsampled). As can be seen, although some of the components can still be detected for TRs up to 950ms, the ultrafast sampling rate used in this work reveals a larger number of components, clearly defined within cortical and subcortical structures.

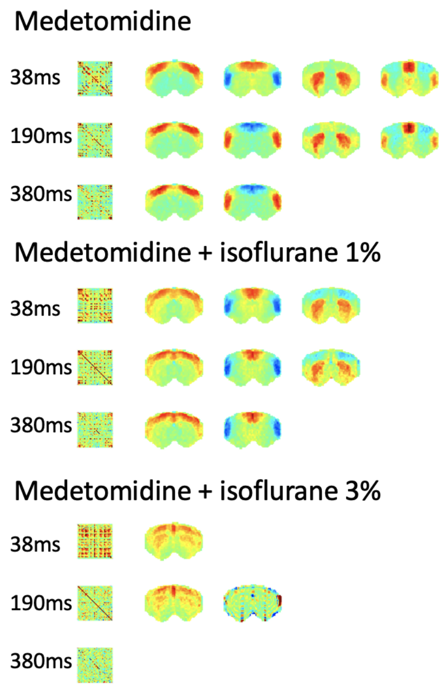


**Fig. S13 - Eigenvectors of the correlation matrix with eigenvalue above postmortem baseline obtained for different sampling rates.** Before the analysis, the fMRI signals recorded with a TR of 38 milliseconds (ms) were downsampled by considering only one in every 5, 10 and 15frames (corresponding to intervals of 190 and 380 milliseconds between frames). The number of eigenvectors detected corresponds to the number of eigenvalues larger than the largest eigenvalue from a postmortem scan (signals also downsampled). As can be seen, although similar components can be detected using the correlation instead of the covariance matrix, the correlation reveals less components, with no component being detected in the deep anesthesia condition for a TR of 380 ms.

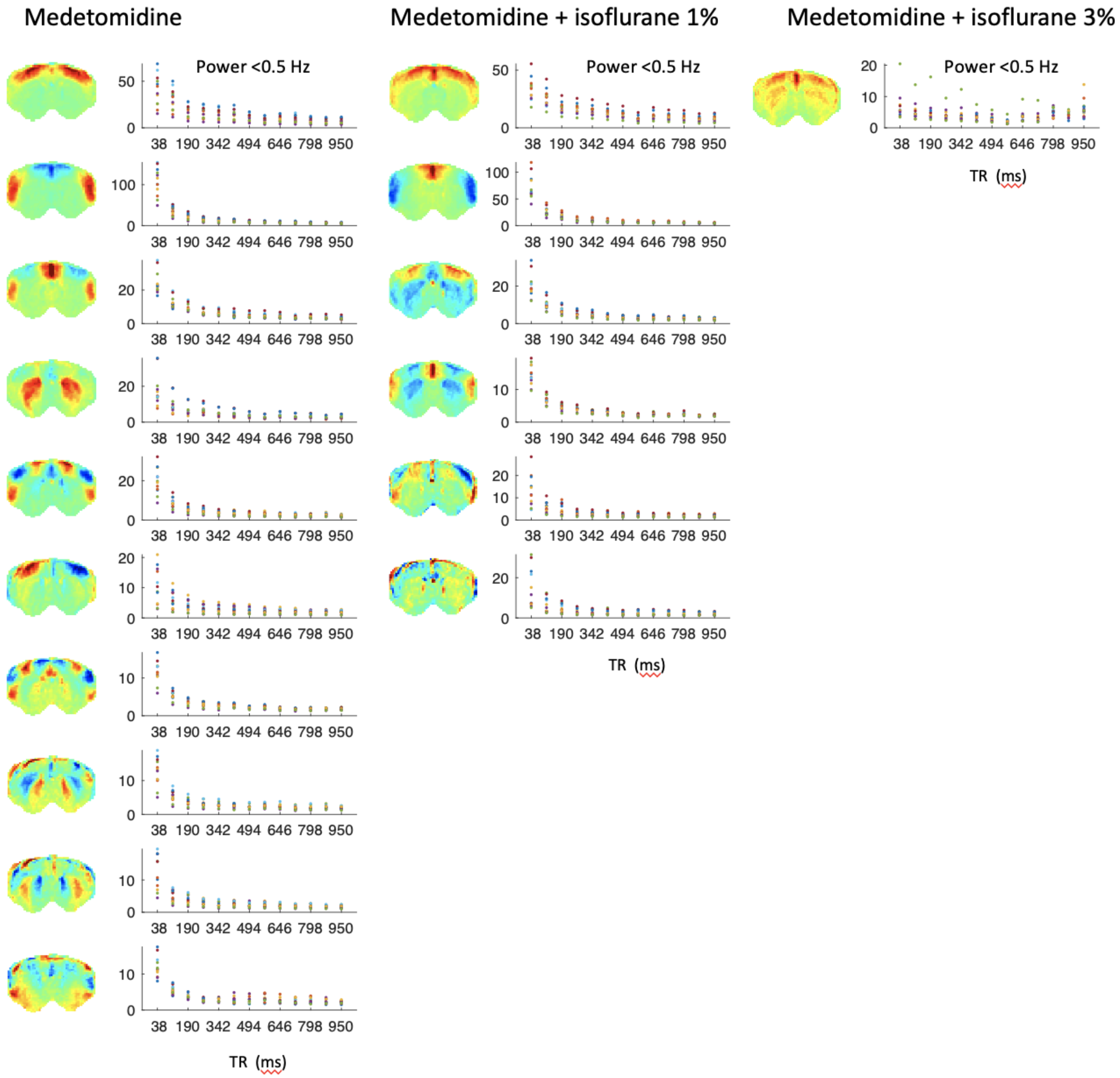


**Fig. S14 - Effect of the sampling rate on the power at low frequencies with respect to postmortem baseline.** Before the analysis, the fMRI signals recorded with a TR of 38 milliseconds (ms) were downsampled by considering only one in every 5, 10, 15, 25 frames (corresponding to intervals of 190 to 950 milliseconds between frames). Subsequently, considering the spatial maps of the principal components detected across conditions at the original ultrafast sampling, the associated temporal signatures are obtained for each scan and the power spectra computed. To correct for the fact that sparser sampling also reduces the number of observed time points, the power is normalized by the average power in the 2 postmortem scans downsampled at the same rates. As can be seen, given that less points per scan are recorded, the power at low frequencies decreases sharply as the sampling rate increases.

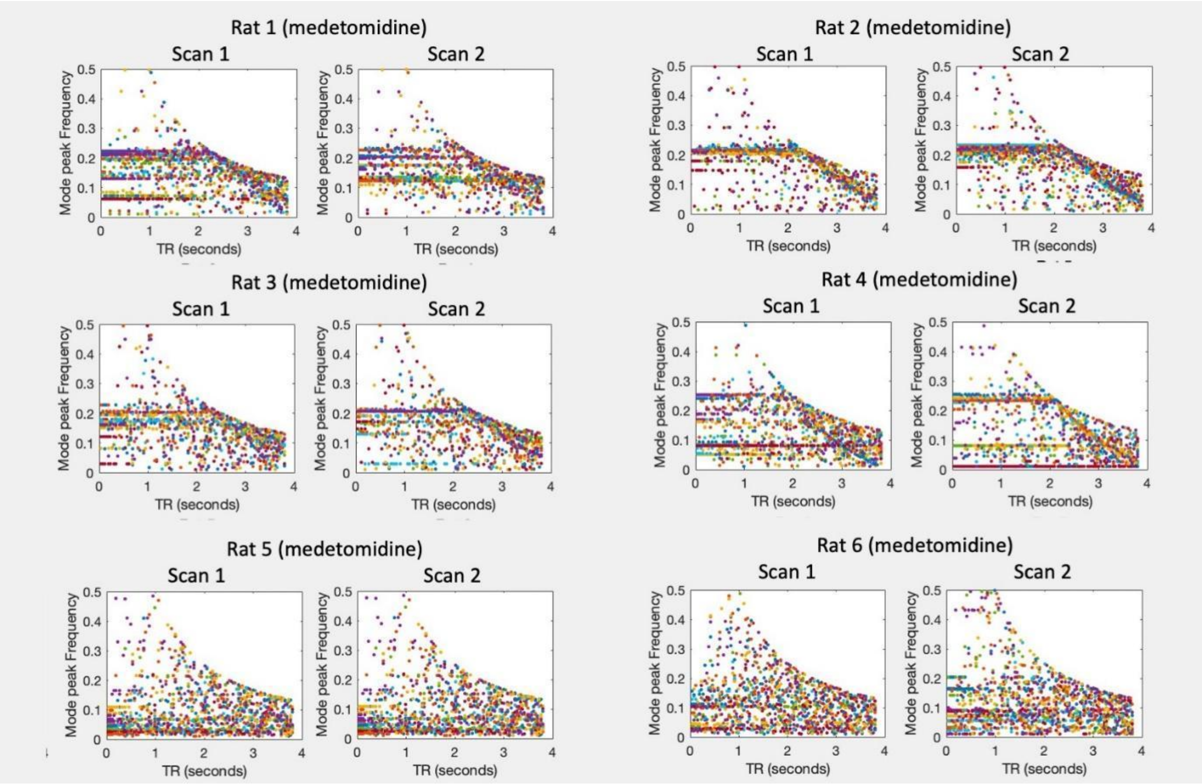


**Fig. S15. Peak frequencies associated with the 10 principal components (color coded) as a function of the temporal resolution.** TR = Repetition Time of fMRI acquisitions. The ultrafast fMRI data was downsampled by considering , the temporal signature associated to each of the 10 covariance modes shown in Figure 3 in the manuscript was obtained, and the peak frequency of the associated power spectrum is reported, for each scan of sedated rats. Horizontal lines in the plots mean that the peak frequency can be detected at different temporal resolutions, with some rats showing robustness up to the Nyquist frequency. Note that, for a TR = 2s, the fastest frequency detected is 0.25 Hz, such that for longer TRs, only resonance at slower frequencies can be detected.

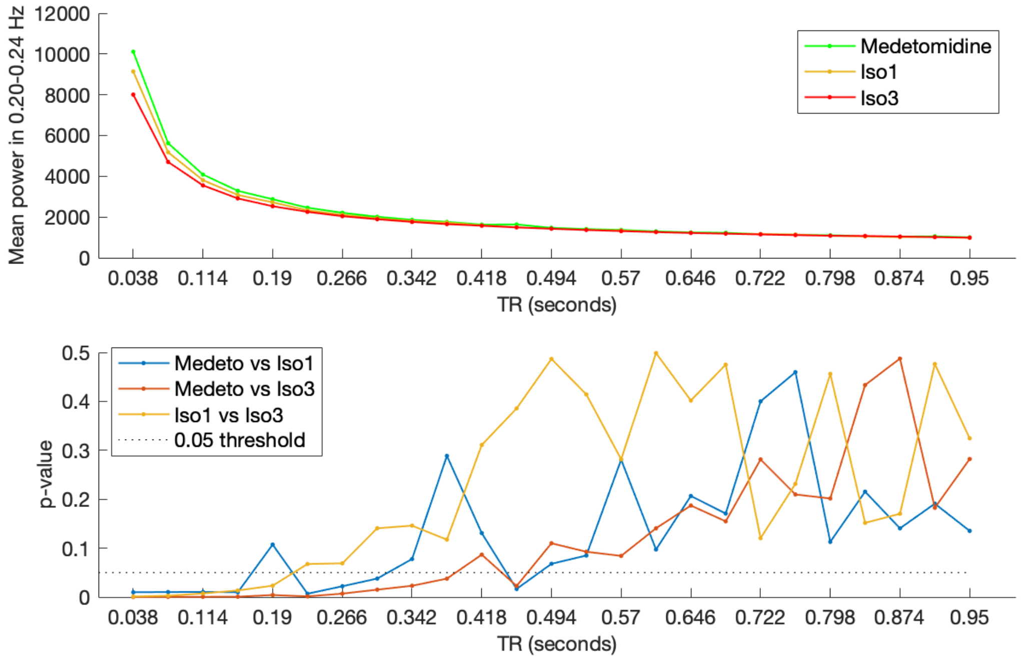


**Fig. S16. High temporal resolution improves the detection of differences in power at specific frequencies between conditions.** To investigate the effects of the temporal resolution, the fMRI signals were downsampled at reduced temporal resolution to study the impact on the results. (Top) The power at 0.2-0.24 Hz (where most significant differences were detected between conditions) was found to substantially decrease as the sampling rate was decreased. (Bottom) The differences detected between conditions were found to lose significance (p-values >0.05) for TR > 0.152 seconds (bottom plot). TR = repetition time of fMRI acquisitions reported in seconds.

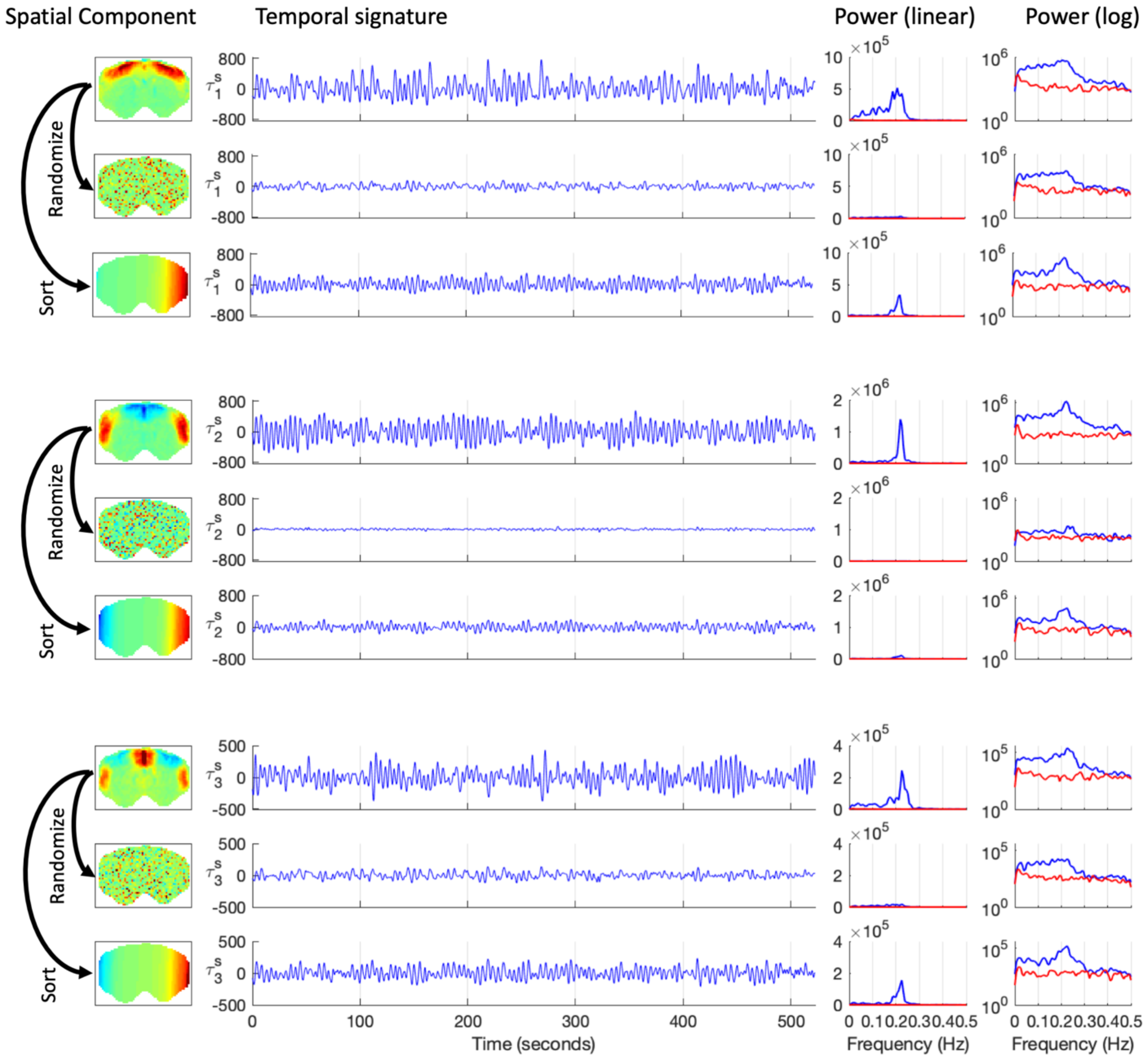


**Fig. S17 - The oscillations are specific to the spatial patterns.** To demonstrate that the oscillations are specific to the spatial wave patterns detected, we reorder the elements in 3 representative spatial patterns by randomizing the voxels or sorting them according to the phase relationships, ensuring a spatial gradient. This analysis demonstrates that, when projecting the *randomized* spatial patterns (with size 1xN) to the same, unfiltered, fMRI signals (with size NxT), no oscillations are detected in the associated temporal signature (with size 1xT), with the power being close to what is detected from a postmortem scan (red lines). The power spectra are shown in both linear and logarithmic (log) scales. When the elements are sorted to keep a spatial gradient (increasing from left to right), we still find oscillations, but with lower amplitude and uncorrelated with the original temporal signature. This occurs because the sorted patterns approximate a different intrinsic spatial pattern.

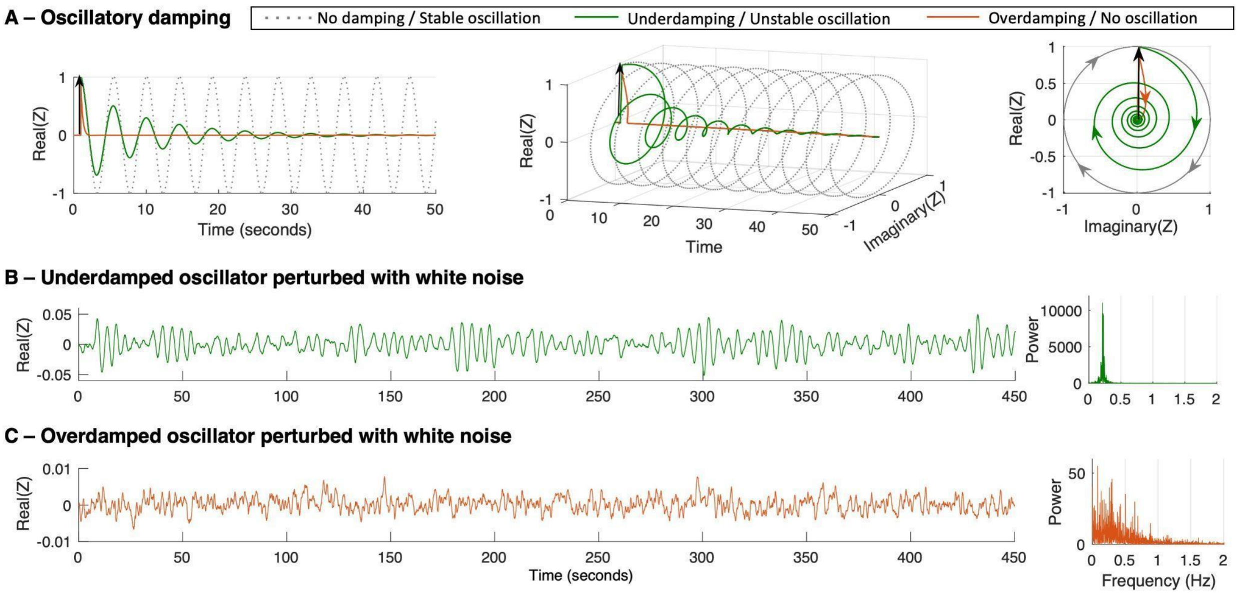


**Fig. S18 – Regimes of oscillatory damping. A** – Illustration of 3 regimes of oscillatory motion through numerical integration of the Stuart Landau equation in response to a unit pulse added at t = 1 s with $\omega_0$ = 0.22 Hz and 3 distinct levels of damping: $a$ = 1 (no damping, dashed gray line), $a$ = -0.1 (underdamping, green line) and $a$ =-2 (overdamping, red line). Since Z has an imaginary component – ensuring the conservation of angular momentum - 3 perspectives of the same plot are reported in left, middle and right panels. **B,C** – Simulation of a Hopf oscillator with natural frequency $\omega_0$ = 0.22 Hz in the underdamped (B) and overdamped (C) regimes in the presence of Gaussian white noise and corresponding power spectra.

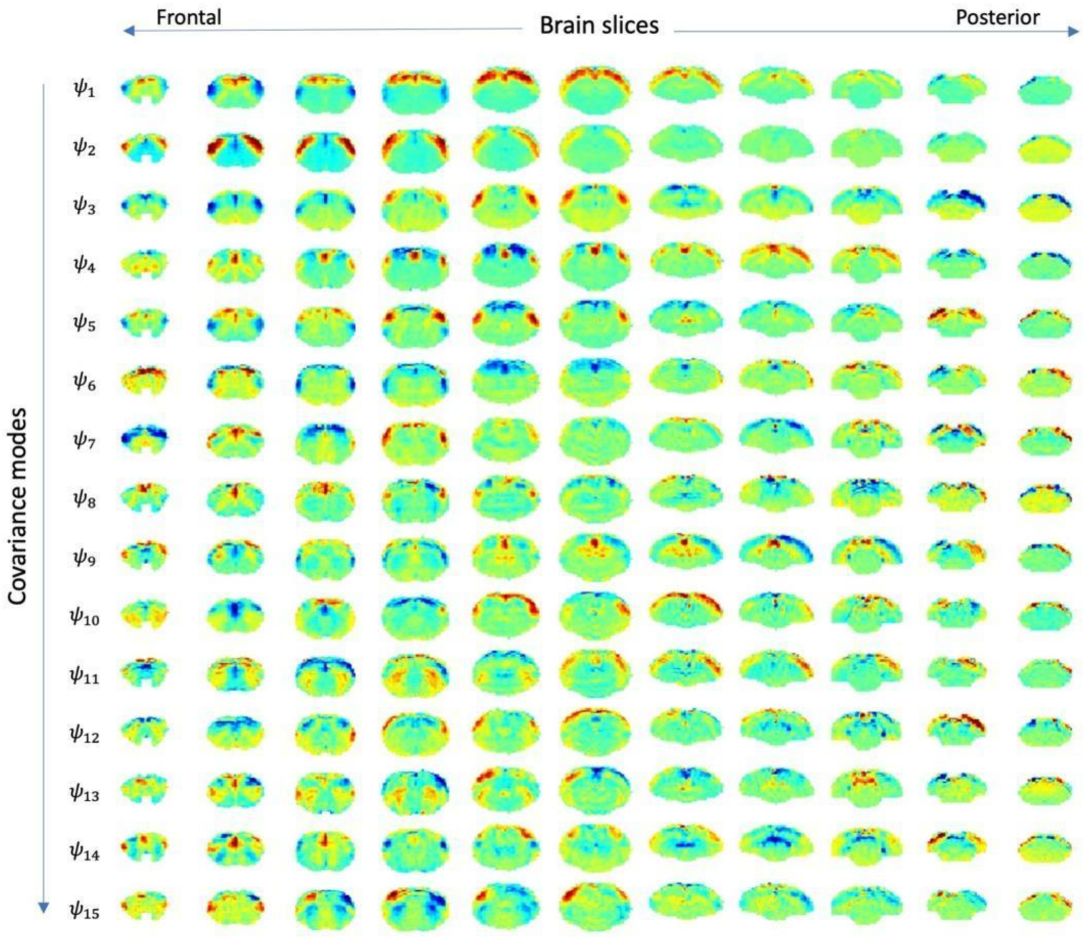


**Fig. S19. Covariance modes detected from 6 whole-brain fMRI scans.** The 15 eigenvectors with largest magnitude eigenvalue obtained from the covariance matrix of fMRI signals band-pass filtered between 0.05 and 0.25 Hz averaged across the 6 whole-brain scans from 3 sedated rats under medetomidine.

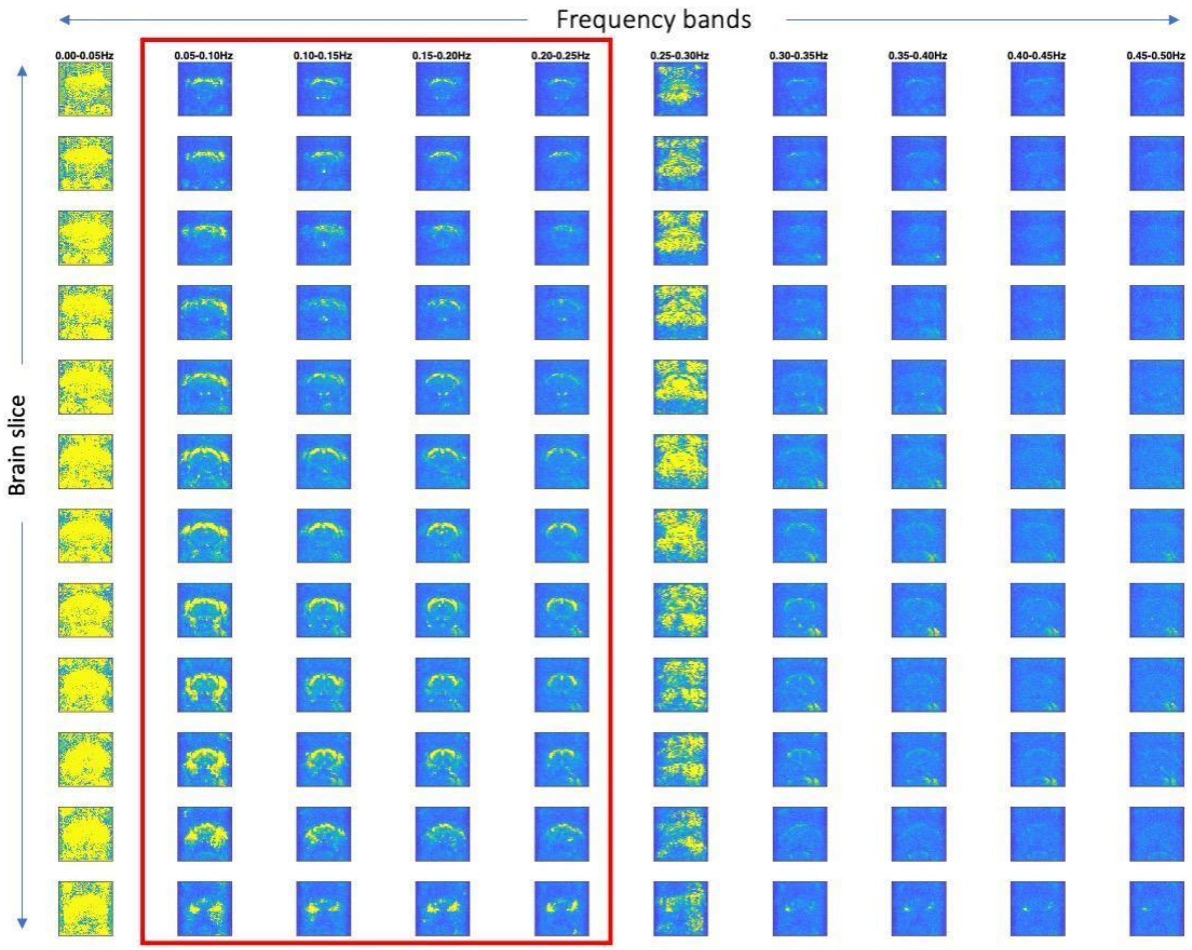


**Fig. S20. Space-Frequency analysis of an individual whole-brain fMRI scan in a sub-range of frequencies.** From the same fMRI scan shown in Fig. S24, voxels in each slice are colored according to the absolute power in each of 10 non-overlapping 0.05Hz-wide frequency bands, revealing that signal artifacts fall below 0.0.5 Hz and between 0.25 and 0.30 Hz, such that an artifact-free frequency range, was detected between 0.05 and 0.25 Hz, revealing similar patterns as the ones detected in single-slice analysis. The artifacts detected in the 1st and 5th bands are consistent across the 6 whole-brain fMRI scans (shown on request).

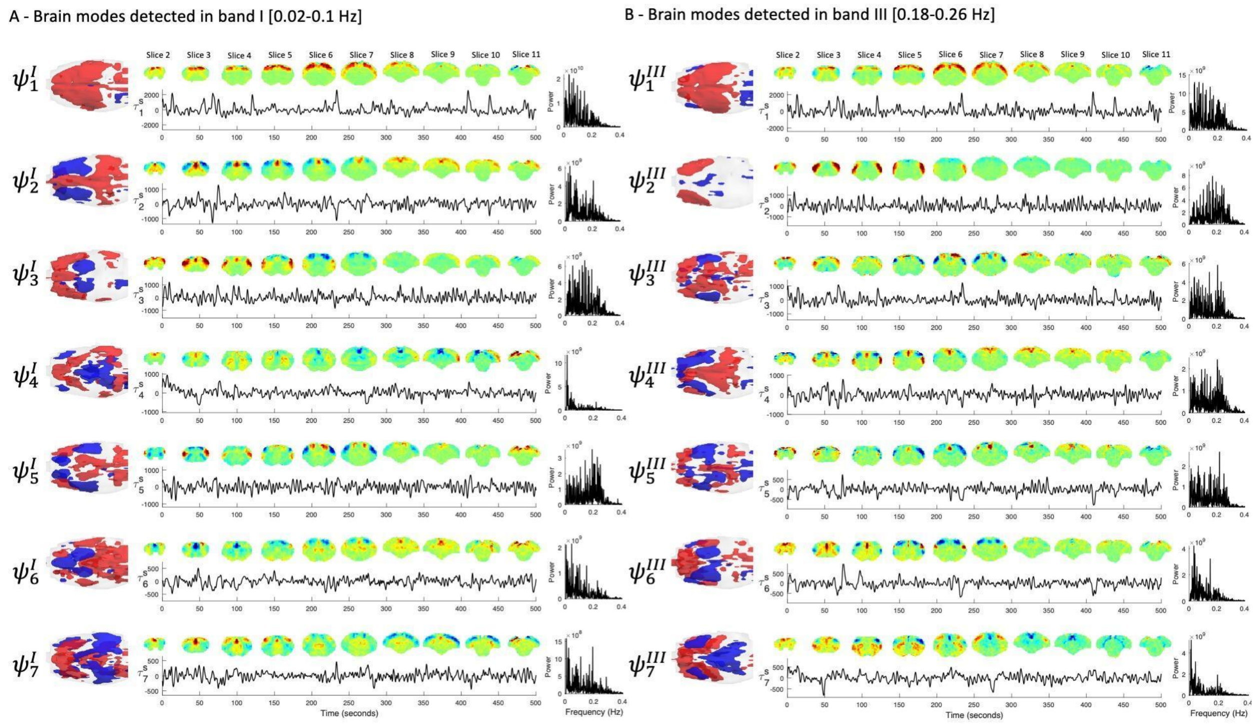


**Fig. S21. Lower temporal resolution of whole-brain scans blurs the detection of resonant oscillations.** The first 7 principal components detected in two different frequency bands (left) 0.02-0.1 Hz, falling in the range typically considered in resting-state fMRI studies and (right) 0.18-0.26 Hz, where the resonant frequency of most modes was detected in the single-slice analysis, from 2 whole brain scans of the same sedated rat (TR = 350ms). For each covariance mode, the spatial configuration is represented in each individual slice and in a 3-dimensional rendering, coloring in red/blue the eigenvector elements above/below 1 standard deviation. In addition, the temporal signature obtained from one scan and the corresponding power spectra are reported for each mode.

| | Rat | Condition | Isoflurane (%) | Time after induction (minutes) | Breathing (Hz) | | | Temperature (°C) | | |
|---|---|---|---|---|---|---|---|---|---|---|
| | | | | | min | max | mean | min | max | mean |
| Frontal Slice | 1 | Sedation | 0 | 113 | 1.72 | 1.75 | 1.73 | 38.2 | 38.4 | 38.3 |
| | | | | 128 | 1.75 | 1.83 | 1.79 | 38.2 | 38.2 | 38.2 |
| | | Light Anaesthesia | 1 | 156 | 1.20 | 1.28 | 1.24 | 38.4 | 38.4 | 38.4 |
| | | | | 170 | 1.17 | 1.23 | 1.20 | 38.3 | 38.4 | 38.4 |
| | | Deep Anaesthesia | 3 | 196 | 0.80 | 0.83 | 0.82 | 38.6 | 38.7 | 38.7 |
| | | | | 211 | 0.80 | 0.83 | 0.82 | 38.5 | 38.6 | 38.6 |
| | 2 | Sedation | 0 | 100 | 1.93 | 2.22 | 2.08 | 37.0 | 37.2 | 37.1 |
| | | | | 115 | 2.00 | 2.08 | 2.04 | 37.2 | 37.3 | 37.3 |
| | | Light Anaesthesia | 1 | 143 | 1.30 | 1.42 | 1.36 | 37.5 | 37.5 | 37.5 |
| | | | | 156 | 1.33 | 1.40 | 1.37 | 37.4 | 37.4 | 37.4 |
| | | Deep Anaesthesia | 3 | 183 | 0.83 | 0.90 | 0.87 | 38.0 | 38.1 | 38.1 |
| | | | | 197 | 0.83 | 0.87 | 0.85 | 38.1 | 38.2 | 38.2 |
| | 3 | Sedation | 0 | 107 | 1.40 | 1.45 | 1.43 | 37.0 | 37.1 | 37.1 |
| | | | | 120 | 1.40 | 1.47 | 1.43 | 36.9 | 37.0 | 37.0 |
| | | Light Anaesthesia | 1 | 147 | 0.87 | 0.97 | 0.92 | 36.6 | 36.8 | 36.7 |
| | | | | 160 | 0.80 | 0.87 | 0.83 | 36.3 | 36.5 | 36.4 |
| | | Deep Anaesthesia | 3 | 188 | 0.75 | 0.80 | 0.78 | 37.1 | 37.3 | 37.2 |
| | | | | 201 | 0.78 | 0.80 | 0.79 | 37.2 | 37.2 | 37.2 |
| | 4 | Sedation | 0 | 158 | 1.92 | 2.07 | 1.99 | 38.9 | 38.9 | 38.9 |
| | | | | 171 | 1.93 | 2.07 | 2.00 | 38.9 | 39.0 | 39.0 |
| | | Light Anaesthesia | 1 | 199 | 1.47 | 1.63 | 1.55 | 38.6 | 39.0 | 38.8 |
| | | | | 211 | 1.47 | 1.52 | 1.49 | 38.3 | 38.5 | 38.4 |
| | | Deep Anaesthesia | 3 | 238 | 0.78 | 0.87 | 0.83 | 38.0 | 38.3 | 38.2 |
| | | | | 251 | 0.72 | 0.78 | 0.75 | 38.0 | 38.0 | 38.0 |
| | 5 | Sedation | 0 | 101 | 1.55 | 1.62 | 1.58 | 36.9 | 37.3 | 37.1 |
| | | | | 114 | 1.45 | 1.50 | 1.48 | 37.5 | 37.7 | 37.6 |
| | | Light Anaesthesia | 1 | 141 | 0.98 | 1.10 | 1.04 | 37.6 | 37.8 | 37.7 |
| | | | | 154 | 0.97 | 1.02 | 0.99 | 37.4 | 37.5 | 37.5 |
| | | Deep Anaesthesia | 3 | 181 | 0.63 | 0.67 | 0.65 | 37.3 | 37.5 | 37.4 |
| | | | | 193 | 0.63 | 0.65 | 0.64 | 37.1 | 37.3 | 37.2 |
| | 6 | Sedation | 0 | 94 | 1.95 | 2.08 | 2.02 | 37.1 | 37.3 | 37.2 |
| | | | | 106 | 1.83 | 1.92 | 1.88 | 37.3 | 37.3 | 37.3 |
| | | Light Anaesthesia | 1 | 134 | 1.18 | 1.42 | 1.30 | 36.8 | 37.2 | 37.0 |
| | | | | 146 | 1.15 | 1.25 | 1.20 | 36.6 | 36.7 | 36.7 |
| | | Deep Anaesthesia | 3 | 174 | 0.77 | 0.85 | 0.81 | 36.4 | 36.8 | 36.6 |
| | | | | 186 | 0.75 | 0.78 | 0.77 | 36.3 | 36.4 | 36.4 |
| Whole brain | 4 | Sedation | 0 | 114 | 1.77 | 1.88 | 1.83 | 36.5 | 37.3 | 36.9 |
| | | | | 132 | 1.85 | 1.95 | 1.90 | 37.5 | 38.5 | 38.0 |
| | 5 | Sedation | 0 | 60 | 1.38 | 1.43 | 1.41 | 35.8 | 36.4 | 36.1 |
| | | | | 79 | 1.42 | 1.62 | 1.52 | 35.7 | 36.4 | 36.1 |
| | 6 | Sedation | 0 | 55 | 1.67 | 1.83 | 1.75 | 36.1 | 36.5 | 36.3 |
| | | | | 74 | 1.70 | 1.90 | 1.80 | 36.2 | 36.8 | 36.5 |

**Table S1. Animal preparation details and physiological monitoring during each scan.** n.a. = not applicable; min = minimum, max = maximum.

A - Statistical differences in peak frequency (p-values)

| | | Sedation | | | | | | | | | Light Anaesthesia | | | | | | Deep An. |
|---|---|---|---|---|---|---|---|---|---|---|---|---|---|---|---|---|---|
| | Mode | 2 | 3 | 4 | 5 | 6 | 7 | 8 | 9 | 10 | 1 | 2 | 3 | 4 | 5 | 6 | 1 |
| Sedation | 1 | 9.0E-02 | 4.9E-01 | 1.1E-01 | 3.2E-01 | 4.6E-01 | 6.5E-02 | 1.9E-01 | 4.6E-01 | 4.3E-01 | 2.3E-01 | 6.8E-02 | 2.2E-02 | 9.5E-02 | 3.6E-01 | 1.4E-01 | 2.3E-05 |
| | 2 | | 8.3E-02 | 1.2E-02 | 2.6E-01 | 7.0E-02 | 4.6E-01 | 1.2E-02 | 4.5E-02 | 1.5E-01 | 7.1E-03 | 5.8E-04 | 7.1E-04 | 3.4E-05 | 1.9E-02 | 1.0E-02 | 1.2E-05 |
| | 3 | | | 1.0E-01 | 2.4E-01 | 4.7E-01 | 9.1E-02 | 2.0E-01 | 4.6E-01 | 3.8E-01 | 3.1E-01 | 1.0E-01 | 6.2E-02 | 8.0E-02 | 3.5E-01 | 1.1E-01 | 1.3E-05 |
| | 4 | | | | 6.7E-02 | 1.3E-01 | 3.8E-03 | 3.6E-01 | 1.8E-01 | 8.0E-02 | 1.5E-01 | 3.7E-01 | 3.4E-01 | 2.7E-01 | 3.2E-01 | 4.9E-01 | 2.6E-05 |
| | 5 | | | | | 3.4E-01 | 2.7E-01 | 9.8E-02 | 1.6E-01 | 3.2E-01 | 1.2E-01 | 5.4E-03 | 6.9E-03 | 3.5E-02 | 1.6E-01 | 6.6E-02 | 1.2E-05 |
| | 6 | | | | | | 1.1E-01 | 2.0E-01 | 4.1E-01 | 5.0E-01 | 2.6E-01 | 9.0E-02 | 6.5E-02 | 4.3E-02 | 2.3E-01 | 1.2E-01 | 2.0E-05 |
| | 7 | | | | | | | 2.3E-02 | 4.8E-02 | 1.1E-01 | 2.8E-02 | 7.4E-03 | 4.2E-03 | 9.0E-03 | 4.1E-02 | 3.3E-03 | 1.8E-05 |
| | 8 | | | | | | | | 2.2E-01 | 1.2E-01 | 2.6E-01 | 2.5E-01 | 3.0E-01 | 3.1E-01 | 3.1E-01 | 4.7E-01 | 1.9E-05 |
| | 9 | | | | | | | | | 3.9E-01 | 3.3E-01 | 1.6E-01 | 8.3E-02 | 8.5E-02 | 4.1E-01 | 1.9E-01 | 1.3E-05 |
| | 10 | | | | | | | | | | 2.5E-01 | 2.7E-02 | 2.9E-02 | 8.5E-02 | 3.2E-01 | 9.1E-02 | 1.3E-05 |
| Light Anaesthesia | 1 | | | | | | | | | | | 6.0E-02 | 3.5E-02 | 6.6E-02 | 5.0E-01 | 2.0E-01 | 9.0E-06 |
| | 2 | | | | | | | | | | | | 4.6E-01 | 4.2E-01 | 1.2E-01 | 3.5E-01 | 1.2E-05 |
| | 3 | | | | | | | | | | | | | 4.7E-01 | 9.9E-02 | 3.3E-01 | 7.8E-06 |
| | 4 | | | | | | | | | | | | | | 1.5E-01 | 3.0E-01 | 1.1E-05 |
| | 5 | | | | | | | | | | | | | | | 2.5E-01 | 2.3E-05 |
| | 6 | | | | | | | | | | | | | | | | 1.5E-05 |

B - A - Statistical differences in Q-factor (p-values)

| | | Sedation | | | | | | | | | Light Anaesthesia | | | | | | Deep An. |
|---|---|---|---|---|---|---|---|---|---|---|---|---|---|---|---|---|---|
| | Mode | 2 | 3 | 4 | 5 | 6 | 7 | 8 | 9 | 10 | 1 | 2 | 3 | 4 | 5 | 6 | 1 |
| Sedation | 1 | 9.0E-02 | 4.9E-01 | 1.1E-01 | 3.2E-01 | 4.6E-01 | 6.5E-02 | 1.9E-01 | 4.6E-01 | 4.3E-01 | 2.3E-01 | 6.8E-02 | 2.2E-02 | 9.5E-02 | 3.6E-01 | 1.4E-01 | 2.3E-05 |
| | 2 | | 8.3E-02 | 1.2E-02 | 2.6E-01 | 7.0E-02 | 4.6E-01 | 1.2E-02 | 4.5E-02 | 1.5E-01 | 7.1E-03 | 5.8E-04 | 7.1E-04 | 3.4E-05 | 1.9E-02 | 1.0E-02 | 1.2E-05 |
| | 3 | | | 1.0E-01 | 2.4E-01 | 4.7E-01 | 9.1E-02 | 2.0E-01 | 4.6E-01 | 3.8E-01 | 3.1E-01 | 1.0E-01 | 6.2E-02 | 8.0E-02 | 3.5E-01 | 1.1E-01 | 1.3E-05 |
| | 4 | | | | 6.7E-02 | 1.3E-01 | 3.8E-03 | 3.6E-01 | 1.8E-01 | 8.0E-02 | 1.5E-01 | 3.7E-01 | 3.4E-01 | 2.7E-01 | 3.2E-01 | 4.9E-01 | 2.6E-05 |
| | 5 | | | | | 3.4E-01 | 2.7E-01 | 9.8E-02 | 1.6E-01 | 3.2E-01 | 1.2E-01 | 5.4E-03 | 6.9E-03 | 3.5E-02 | 1.6E-01 | 6.6E-02 | 1.2E-05 |
| | 6 | | | | | | 1.1E-01 | 2.0E-01 | 4.1E-01 | 5.0E-01 | 2.6E-01 | 9.0E-02 | 6.5E-02 | 4.3E-02 | 2.3E-01 | 1.2E-01 | 2.0E-05 |
| | 7 | | | | | | | 2.3E-02 | 4.8E-02 | 1.1E-01 | 2.8E-02 | 7.4E-03 | 4.2E-03 | 9.0E-03 | 4.1E-02 | 3.3E-03 | 1.8E-05 |
| | 8 | | | | | | | | 2.2E-01 | 1.2E-01 | 2.6E-01 | 2.5E-01 | 3.0E-01 | 3.1E-01 | 3.1E-01 | 4.7E-01 | 1.9E-05 |
| | 9 | | | | | | | | | 3.9E-01 | 3.3E-01 | 1.6E-01 | 8.3E-02 | 8.5E-02 | 4.1E-01 | 1.9E-01 | 1.3E-05 |
| | 10 | | | | | | | | | | 2.5E-01 | 2.7E-02 | 2.9E-02 | 8.5E-02 | 3.2E-01 | 9.1E-02 | 1.3E-05 |
| Light Anaesthesia | 1 | | | | | | | | | | | 6.0E-02 | 3.5E-02 | 6.6E-02 | 5.0E-01 | 2.0E-01 | 9.0E-06 |
| | 2 | | | | | | | | | | | | 4.6E-01 | 4.2E-01 | 1.2E-01 | 3.5E-01 | 1.2E-05 |
| | 3 | | | | | | | | | | | | | 4.7E-01 | 9.9E-02 | 3.3E-01 | 7.8E-06 |
| | 4 | | | | | | | | | | | | | | 1.5E-01 | 3.0E-01 | 1.1E-05 |
| | 5 | | | | | | | | | | | | | | | 2.5E-01 | 2.3E-05 |
| | 6 | | | | | | | | | | | | | | | | 1.5E-05 |

**Table S2. Statistical comparison of mode Q-factors between conditions.** p-values from a permutation test with 1000 permutations. yellow: p-value <0.05; red: p-value < 0.05/136 (Bonferroni).
